# Memorability of photographs in subjective cognitive decline and mild cognitive impairment: implications for cognitive assessment

**DOI:** 10.1101/660365

**Authors:** Wilma A. Bainbridge, David Berron, Hartmut Schütze, Arturo Cardenas-Blanco, Coraline Metzger, Laura Dobisch, Daniel Bittner, Wenzel Glanz, Annika Spottke, Janna Rudolph, Frederic Brosseron, Katharina Buerger, Daniel Janowitz, Klaus Fliessbach, Michael Heneka, Christoph Laske, Martina Buchmann, Oliver Peters, Dominik Diesing, Siyao Li, Josef Priller, Eike Jakob Spruth, Slawek Altenstein, Anja Schneider, Barbara Kofler, Stefan Teipel, Ingo Kilimann, Jens Wiltfang, Claudia Bartels, Steffen Wolfsgruber, Michael Wagner, Frank Jessen, Chris Baker, Emrah Düzel

## Abstract

**INTRODUCTION:** Impaired long-term memory is a defining feature of Mild Cognitive Impairment (MCI). We tested whether this impairment is item-specific, limited to some memoranda whereas some remain consistently memorable.

**METHODS:** We conducted item-based analyses of long-term visual recognition memory. 394 participants (healthy controls (HC), Subjective Cognitive Decline (SCD), and MCI) in the multicentric DZNE-Longitudinal Cognitive Impairment and Dementia Study (DELCODE) were tested with images from a pool of 835 photographs.

**RESULTS:** We observed consistent memorability for images in HCs, SCDs, and MCI, predictable by a neural network trained on another healthy sample. Looking at memorability differences between groups, we identified images that could successfully categorize group membership with higher success and a substantial image reduction than the original image set.

**DISCUSSION:** Individuals with SCD and MCI show consistent memorability for specific items, while other items show significant diagnosticity. Certain stimulus features could optimize diagnostic assessment, while others could support memory.

## 1. Background

Recent work in healthy individuals has found that certain images are intrinsically memorable or forgettable across observers [1,2]; there are images of faces or scenes that most people remember or forget, regardless of their different individual experiences. This *memorability* of an image can be quantified and predicts 50% of the variance in people’s performance on a memory test [2]. Viewing memorable images automatically elicits specific neural signatures [3,4], and the memorability score of an image can be predicted by computational models [5,6]. However, image attributes such as aesthetics, emotionality, typicality, or what people believe will be memorable do not fully predict memorability [2,7], and memorability is an automatically processed image property that is resilient to the effects of attention [8]. This means that researchers can predict in advance what images a person is likely to remember or forget, and use such information to create memorable educational materials, or design well-balanced memory tests.

While memorability has so far been characterized based on healthy participants’ memory behavior, it is unclear if memorability is also consistent in populations with memory impairments at increased risk for Alzheimer’s Disease (AD), such as Mild Cognitive Impairment (MCI) or Subjective Cognitive Decline (SCD) [9]. Consistent memorability in SCD and MCI would enable better prediction of what images are likely to be remembered or forgotten. Furthermore, changes in memorability patterns across disease stages could improve cognitive staging and design of cognitive progression markers. By avoiding highly memorable images, cognitive tests could be made more time efficient and more sensitive. Understanding which stimulus features improve or impair memorability could provide insights into the cognitive processes that are impaired. Furthermore, knowledge about memorability could aid in the design of memorable environments, or allow clinicians to focus on aiding memory for forgettable items.

In the current study, we analyzed the results of a visual recognition memory test in which each participant had to memorize a randomly selected subset of 88 photographs from a pool of 835. This randomization afforded us the possibility to assess memorability unconfounded by systematic effects of stimulus-selection or stimulus-order effects. Using data from 394 individuals, including those with SCD, MCI, and healthy controls (HC), we identified two meaningful sets of images: 1) images that can consistently predict performance of participant groups, and 2) images that reliably differentiate groups.

## 2. Methods

### 2.1 Study design

Visual memory tests were analyzed from the DZNE-Longitudinal Cognitive Impairment and Dementia Study (DELCODE), an observational, longitudinal memory clinic-based study across 10 sites in Germany. Specific details about this study, the visual memory task, and data handling and quality control are reported in Jessen et al. [10] and Düzel et al. [11]. The data analyzed in this study were from the second data release from the DELCODE study comprising of 700 individuals of which 394 participants with complete datasets were analyzed, including 136 participants with SCD, 65 with MCI, and 193 HC. Individuals with SCD and MCI were recruited through referrals and self-referrals, while HC were recruited through public advertisements.

The study protocol was approved by all involved centers’ institutional review boards and ethical committees, and all participants gave written informed consent. DELCODE is retrospectively registered at the German Clinical Trials Register (DRKS00007966), (04/05/2015).

### 2.2 Visual memory test

Participants performed an fMRI scene image encoding and retrieval task [12]. First, while in the fMRI scanner, participants studied 88 novel scene target images (44 indoor and 44 outdoor scenes) and 44 repetitions of two pre-familiarized images (one indoor and one outdoor, 22 times each). All images were 8-bit gray scale, presented on an MR-compatible LCD screen (Medres Optostim), scaled to 1250 × 750 pixel resolution and matched for luminance, with a viewing horizontal half-angle of 10.05° across scanners. Each image was presented for 2500ms (with an optimized jitter for statistical efficiency), and participants categorized them as “indoor” or “outdoor” with a button press. Outside of the scanner after a 70-minute delay, participants completed a recognition memory task with these 88 images and 44 novel foil images (22 indoor and 22 outdoor). Participants indicated their recognition memory with a 5-point scale: 1) *I am sure that this picture is new*, 2) *I think that this picture is new*, 3) *I cannot decide if this picture is new or old*, 4) *I think I saw this picture before*, or 5) *I am sure that I did see this picture before*. Results from the fMRI study are reported in [12].

While each participant was tested on 88 target images and 44 foil images, these images were randomly sampled from a larger set of 835 scene images, allowing us to conduct image-based analyses on a large set of images (see Figure 1 for example images). This randomization allowed us to avoid confounding effects of image selection and image order on memory performance. On average, each image served as a target image for 20.3 HC, 14.3 SCD, and 6.8 MCI individuals.

**Figure 1:**
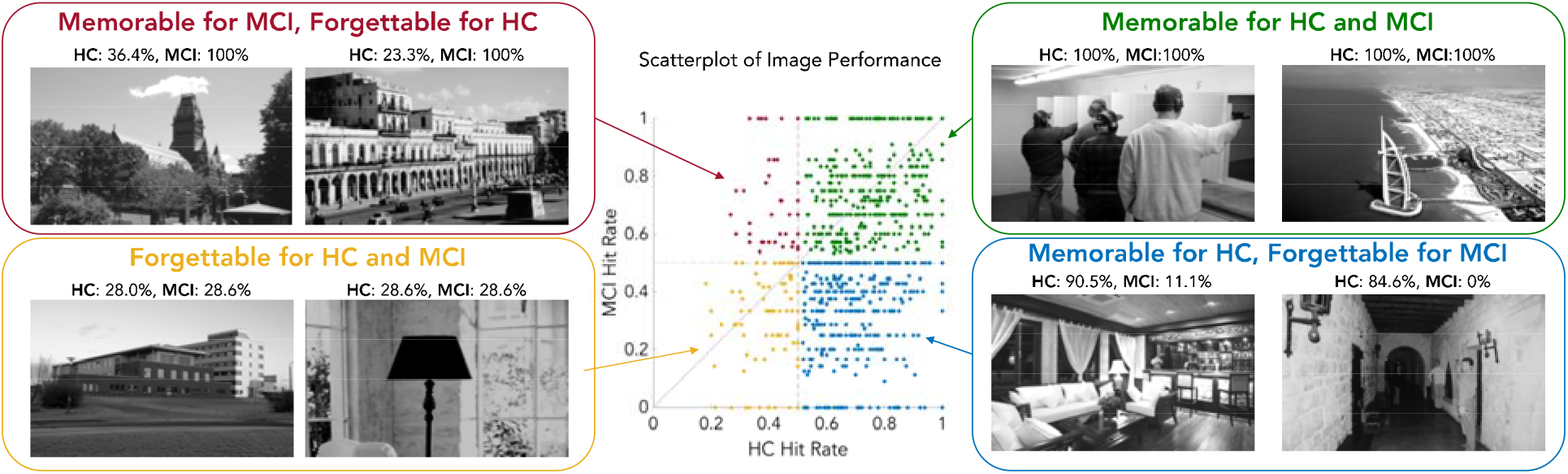
Example images and group performance. The scatterplot shows the distribution of memory performance (hit rate) for all 835 images for healthy controls (HC) versus individuals with Mild Cognitive Impairment (MCI). The diagonal line indicates the points at which performance is equal between both groups. Based on performance, images can be conceptually sorted into four quadrants: 1) images that are memorable to both HC and MCI individuals (green), 2) images that are memorable to HC but forgettable to MCI (blue), 3) images that are forgettable to both groups (yellow), and images that are memorable to MCI but forgettable to HC (red). Example images and performances at the extreme ends for each quadrant are arranged around the scatterplot. In the work that follows, we analyze these four groups of images and determine if they can be used meaningfully to predict memory performance.

### 2.3 Analyzing similarity of MCI, SCD, and healthy individuals: Predicting performance

We first asked whether there are consistencies in memory performance for MCI and SCD just as there are for healthy individuals [1]; i.e., whether there are certain images that patients tend to remember or forget, and, if such consistencies exist, to what degree they align with the images that tend to be remembered and forgotten by HCs.

To address this question, Spearman’s rank correlations of hit rate (HR) performance on images in the visual memory task were calculated between the different patient groups and controls. To assess memorability consistency within patient groups, we conducted a *consistency analysis* as described in Isola et al. [1], where participants are split into random halves (across 1000 iterations) and their hit rates for all images are calculated, and Spearman’s rank correlated between the two halves. We also examined whether a convolutional neural network (CNN) that is significantly able to predict memory performance in healthy individuals [6] could also predict memorability for SCD and MCI groups. MemNet is a CNN with the architecture and pretraining set of Hybrid-CNN [13], a CNN able to classify thousands of object and scene images, then trained to predict the memorability score of an image (i.e., the likelihood for that image to be remembered by any given person). We obtained MemNet scores for each of the 835 stimulus images and used Spearman’s rank correlations to test the degree to which MemNet-predicted memory scores were correlated with patient group memory scores.

### 2.4 Analyzing dissimilarity of MCI, SCD and healthy individuals: Differentiating patient groups

An equally important question is whether there is a set of images in which consistencies in memory performance reliably differ between patient populations and healthy individuals. If such images exist, then they could form an optimized test to distinguish patients from healthy controls with high efficiency.

To explore this question, we conducted an analysis we call the *Iterative Image Subset* (IIS) *Analysis* to compare HC with MCI, and HC with SCD. First, the HC participant pool was randomly downsampled so that the same number of HC were used in the analysis as MCI or SCD individuals. The entire pool of participants was then split into two random halves (Group A and Group B). HR on the memory task was calculated for each image for the HC (*HR*_*GroupA,Healthy*_) and for the patients (*HR*_*GroupA,Patient*_) in Group A. Using this performance metric, we formed three subsets of images. The number of images used in each subset was selected iteratively for all possible subset sizes, ranging from 0% to 100% of images (835 images) in 1% increments, to determine the optimal image subset size. The three resulting subsets were:

1. **“H>P”**, the top set of images where HC outperformed patients (i.e., maximizing *HR*_*GroupA,Healthy*_ - *HR*_*GroupA,Patient*_)
2. **“H<P”**, the top set of images where patients outperformed HC (i.e., maximizing *HR*_*GroupA,Patient*_ - *HR*_*GroupA,Healthy*_)
3. **“H**=**P”**, the top set of images where HC performed most similarly to patients (i.e., minimizing | *HR*_*GroupA,Healthy*_ - *HR*_*GroupA,Patient*_ |)

We then assessed the performance of classifying subjects in Group B using each of the three subsets of images. Specifically, using just the images in a single subset (e.g., H>P), we determined the HR for each of the individuals in Group B (*HR*_*GroupB*_). We then performed a Receiver Operating characteristic (ROC) analysis to determine the diagnostic ability of this subset of images, applying a range of HR cutoffs from 0 to 1 to classify an individual from Group B as either HC or patient, using *HR*_*GroupB*_. We calculated the accuracy of this test based on group membership, and contrasted successful patient diagnosis (true positives) with misclassification of HC (false positives). We assessed classification performance by Area Under the Curve (AUC), where a score of 1 indicates perfect performance, while 0.5 indicates chance performance. This complete analysis was conducted across 100 random participant splits into Group A and B.

### 2.5 Finding image attributes that distinguish these image sets

To see what aspects of the images may determine their membership into different image sets, we conducted an experiment using the online crowd-sourcing platform Amazon Mechanical Turk (AMT). For each of the 835 images, 12 online participants rated the scene in the image on five relevant properties identified in previous scene perception and memorability research [7,14] using a 5-point Likert scale: size (of the portrayed scene), clutter, aesthetics, interest, and whether they think they would remember the image (subjective memorability). They also indicated whether the image showed a natural or manmade scene and if there was a person present. 450 people anonymously participated in the study and provided consent, and this study was approved by the National Institutes of Health (NIH) Office of Human Subjects Research Protections. Two main comparisons were tested for each attribute, using paired samples t-tests: 1) forgettable versus memorable images with similar performance between HC and patients, 2) diagnostic versus non-diagnostic images, where HC and patients differed in their performance. Forgettable and memorable images were identified as the top set of images where both HC and patients had average performance below or above (respectively) median performance, and the difference between groups was minimized (i.e., H=P). Diagnostic and non-diagnostic images were selected from the sets resulting from the IIS analysis (Section 2.4), e.g., H>P and H<P image sets, respectively. The number of images in each set was taken as the optimal number of images identified from the IIS analysis.

## 3. Results

### 3.1 Consistencies in the memories of patient groups

As expected, patient groups of increasing memory impairment showed decreases in average memory performance (HC: M=0.68, SD=0.17; SCD: M=0.62, SD=0.18; MCI: M=0.53, SD=0.26). However, there were also impressive correlations across groups in the images they remembered best or worst (Figure 2). HC and SCD had a significant Spearman’s rank correlation of *ρ*=0.50 (*p*=1.03 × 10^−54^), while HC and MCI had a significant correlation of *ρ*=0.28 (*p*=1.34 × 10^−16^), and SCD and MCI had a significant correlation of *ρ*=0.31 (*p*=2.12 × 10^−19^). HC performance was significantly more similar to SCD performance than MCI performance (*Z*=6.13, *p* ~ 0), and SCD performance was significantly more similar to HC performance than MCI performance (*Z*=5.42, *p* ~ 0). These results indicate that patient groups and healthy elderly individuals tended to remember the same images as each other. All groups were also internally consistent (HC: *ρ*=0.42; SCD: *ρ*=0.32; MCI: *ρ*=0.22; all *p* < 0.0001), meaning a patient will tend to remember similar images to someone else with the same diagnosis.

**Figure 2:**
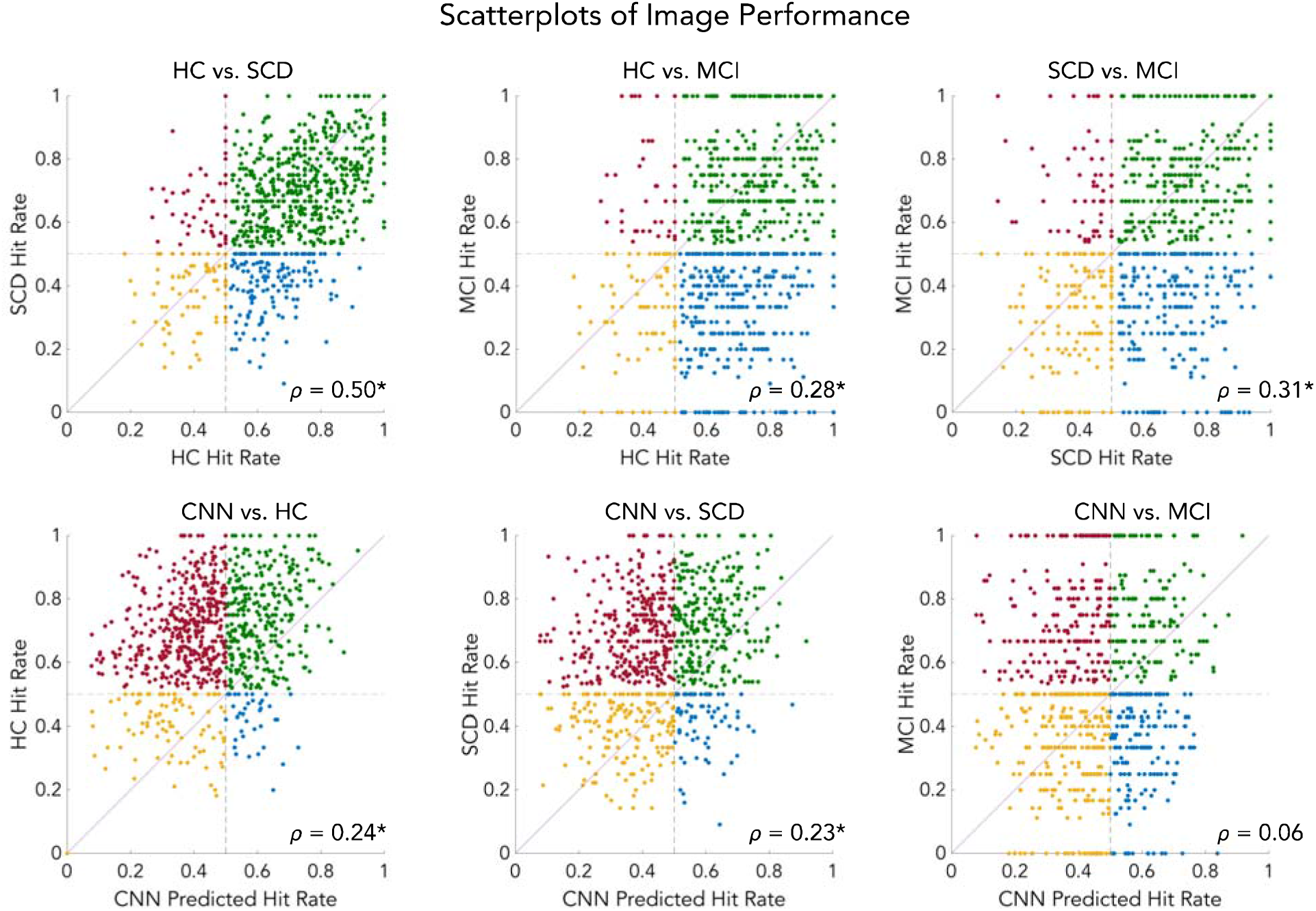
Consistencies across groups and neural networks. The scatterplots show a comparison of hit rates for each of the 835 images between all pairings of the experimental groups (Healthy Controls, HC; Subjective Cognitive Decline, SCD; Mild Cognitive Impairment, MCI), as well as predicted hit rate from a convolutional neural network (CNN) trained to predict memorability scores. Asterisks (*) indicate significant Spearman’s rank correlations. Scatterplot points are colored by quadrant (as in Figure 1), and the diagonal line indicates points where both groups show equal performance.

**Figure 3:**
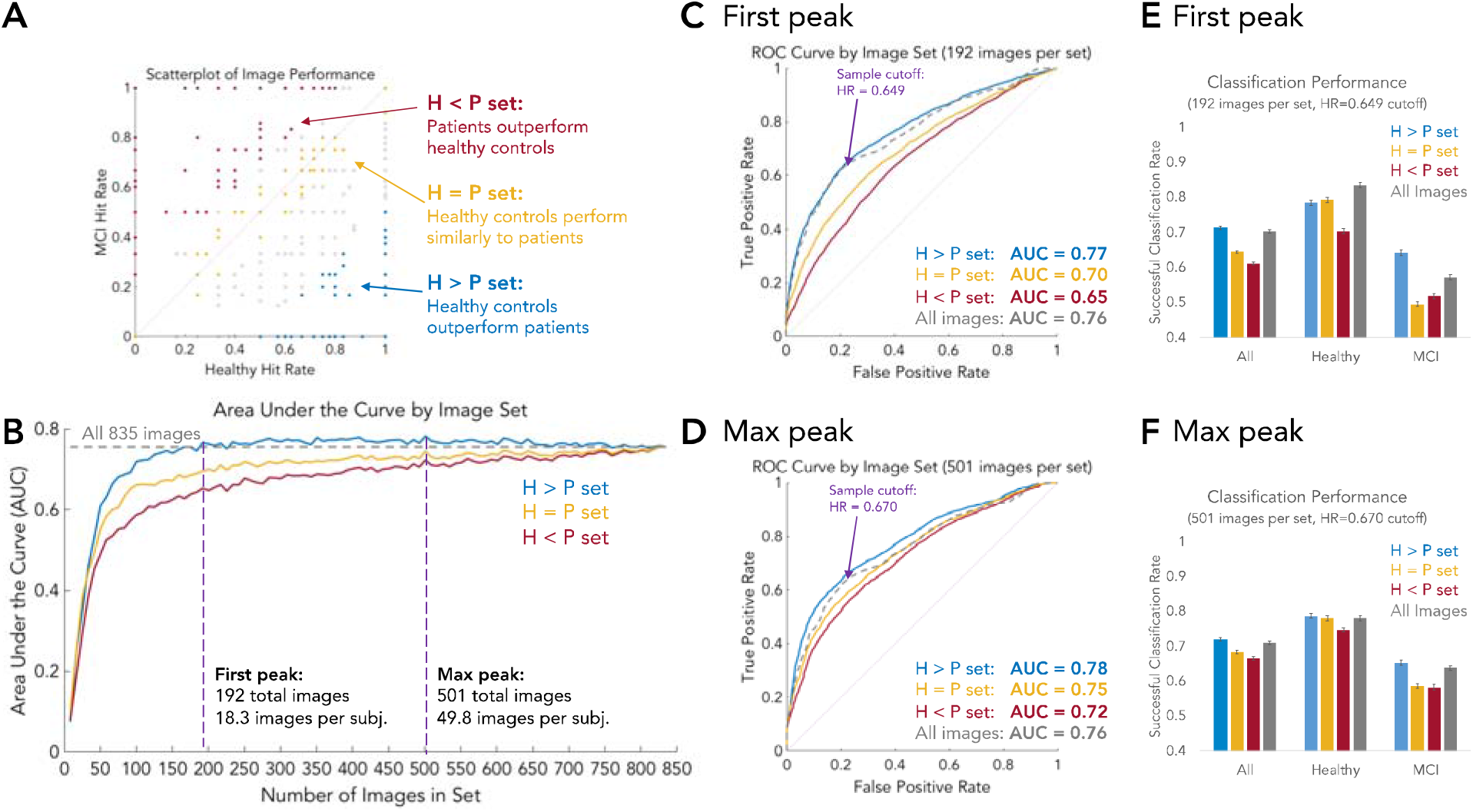
Finding the optimal number of images to diagnose MCI. A) This scatterplot of image performance shows an example of the three possible subsets the images can be divided into: H<P (red), H=P (yellow), and H>P (blue). B) Area Under the Curve (AUC) by image set and number of images in the set. Testing each of these subset types at different set sizes, we find that the H>P set (blue line) consistently outperforms the other image subsets at all set sizes. Importantly, the H>P set also outperforms the all-image set (gray dotted line) at a surprisingly small number of images, first overtaking the all-image set at only 192 images versus the 835 images used in the all-image set. From this set of 192 images, each participant saw on average only 18.3 images. C & D) Receiver Operating Characteristic (ROC) curves for two peaks – the first peak where H>P overtakes the all-image set, and the max peak where H>P has the largest difference from the all-image set. E & F) Participant classification performance, averaged across 100 iterations of participant split-halves, at a sample cutoff (determined as the point where the *true positive rate +* (*1 – false positive rate)* is at its maximum), broken down by participant type for the different image sets. Error bars indicate standard error of the mean across the 100 iterations. Note that the optimized H>P image subset particularly shows a boost in patient diagnosis sensitivity overall other image sets.

The MemNet CNN trained to predict image memorability showed significant correlations with HC (*ρ*=0.24, *p*=3.29 × 10^−12^) and SCD behavior (*ρ*=0.23, *p*=1.84 × 10^−11^), while MCI behavior correlations did not pass significance thresholds (*ρ*=0.06, *p*=0.080).

### 3.2 Differentiating patient groups from healthy controls

As a first test, we examined the ability to differentiate HC and MCI individuals. The IIS analysis shows that the H>P image subset consistently outperforms the H=P and H<P image subsets at all subset sizes, in diagnosing individuals as MCI versus HC (Figure 2). This means that images that are highly memorable to healthy controls but highly forgettable to patients are best able to distinguish these two groups. Surprisingly, H>P image subsets as small as 23% of the original image set were able to surpass the original image set in diagnostic ability. With only 192 total images (or 18.3 images seen per participant), the diagnosis AUC was 0.77, while using the full set of 835 images resulted in an AUC of 0.76. At this 192-image subset size, the difference between subsets is also clear: the H=P set only reaches an AUC of 0.70, while the H<P set performs worse with an AUC of 0.65.

Differentiating HC from SCD individuals shows similar results, even though the two groups have more similar memory performance. The AUC of the H>P set is higher than those of H=P and H<P at all image subset sizes, and the H>P subset first overtakes performance of the full image set at only 92 images in the subset. The AUC for the full image set is 0.59, while with the 92-image subset, the AUC of H>P is also 0.59. In regard to the other image subsets, the AUC for H=P is 0.57, and for H<P it is 0.55. H>P reaches a maximum of performance at a subset size of 367 images, with an AUC of 0.61.

We also determined if the image subsets generalized across groups. We performed the IIS analysis by training on MCI data to determine the image subsets, but then testing those images with SCD data. We find these subsets generalize to each other: the H>P image subset shows higher performance than the other image subsets (H=P, H<P), and first overtakes performance of all images (AUC=0.60) at a subset size of only 100 images (H>P: AUC=0.60; H=P: AUC=0.50; H<P: AUC=0.55). The H>P image subset reaches its peak in performance at 417 images, at an AUC of 0.63.

These results show that using a small, honed subset of images results in higher diagnostic performance than a large, exhaustive set of images, for both SCD and MCI populations. Additionally, using a poor set of images (e.g., H<P) could result in a high diagnosis failure rate. We also find that diagnostic images can successfully transfer across groups; using images that identify MCI can also successfully identify SCD. Since all of the above tests use separate halves of the participants to determine the diagnostic images and to predict group membership, this image diagnosticity is likely to translate to other participant samples as well as other experimental contexts.

### 3.3 Image attributes that distinguish these image sets

Finally, we investigated image attributes related to why an image is memorable to both groups, or why it is diagnostic (Figure 4). Focusing on images that have highly correlated performance between patients and healthy controls, memorable scene images tended to contain more clutter (*t*(191)=2.84, *p*=0.005), appeared more interesting (*t*(191)=3.30, *p*=0.001), and were subjectively more memorable to healthy controls (*t*(191)=3.59, *p*=4.17 × 10^−4^). However, they were not different in scene spatial size (*p*=0.567) nor aesthetics (*p*=0.752). In terms of content, memorable versus forgettable images tended to be manmade rather than natural (forgettable: 76.6% manmade, memorable: 87.0%; *Z*(191)=2.64, *p*=0.008), but were equally likely to be indoors (forgettable: 52.1% indoors; memorable: 50.5%; *p*=0.76) and contain people (forgettable: 7.8% contained people; memorable: 13.0%; *p*=0.09).

**Figure 4:**
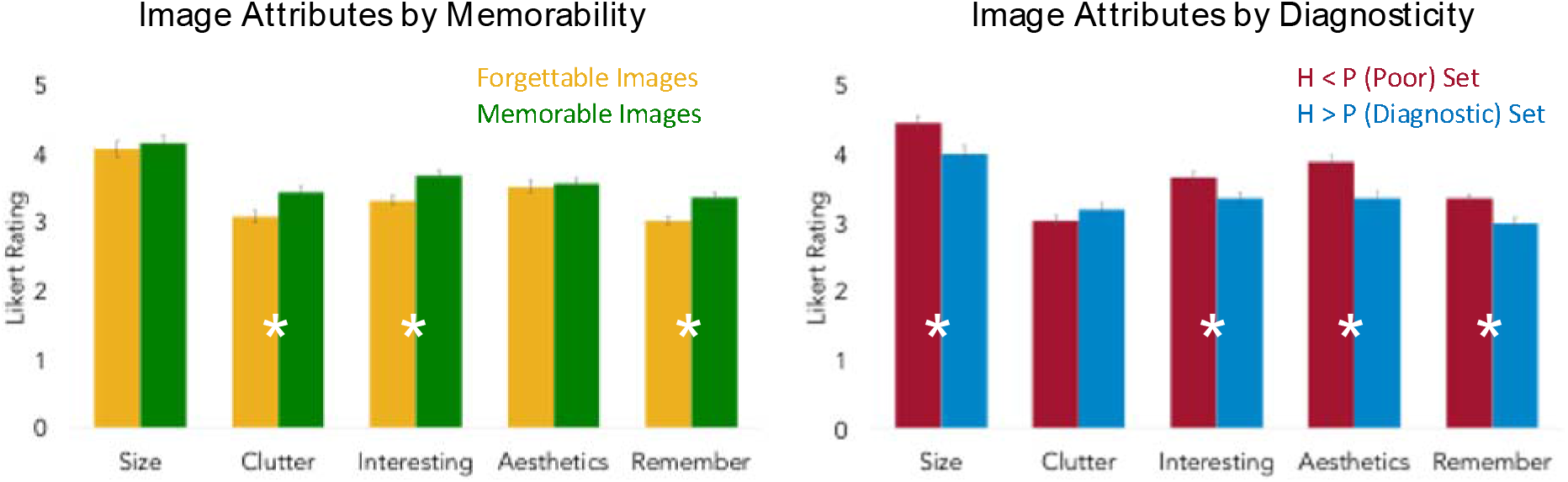
Average attribute ratings based on image set. (Left) Comparison of average attribute ratings between images that are forgettable versus memorable to both HC and individuals with MCI or SCD. (Right) Comparison of average attribute ratings between images from the poorly diagnostic image set (H<P) versus highly diagnostic set (H>P). (Both) All attributes are rated on a Likert scale of 1 (low) to 5 (high). “Remember” is a rating of how likely participants believed they’d be able to remember the image. Asterisks indicate significant differences in a paired samples t-test (p < 0.05). Error bars indicate standard error of the mean.

Focusing on images that show large differences between healthy controls and patients, successfully diagnostic images versus non-diagnostic images tended to be of smaller spaces (*t*(191)=3.05, *p*=0.003), were less interesting (*t*(191)=2.81, *p*=0.005), less aesthetic (*t*(191)=4.04, *p*=7.70 × 10^−5^), and were judged to seem more forgettable by healthy controls (*t*(191)=3.79, *p*=2.05 × 10^−4^), but showed no difference in clutter (*p*=0.153). In terms of content, diagnostic images tended to be manmade (non-diagnostic: 72.4%; diagnostic: 83.9%; *Z*(191)=2.72, *p*=0.007), indoors (non-diagnostic: 37.5%; diagnostic: 55.7%; *Z*(191)=3.58, *p*=3.40 × 10^−4^), and contained people (non-diagnostic: 5.2%; diagnostic: 17.7%; *Z*(191)=3.85, *p*=1.20 × 10^−4^). Memorable images were significantly more interesting (*t*(191)=2.80, *p*=0.006) and seemed subjectively more memorable (*t*(191)=3.55, *p*=4.86 × 10^−4^) than diagnostic images. This shows that diagnostic images that patients forget but healthy controls remember tend to be those that are generally less aesthetic or interesting, yet are manmade, indoor scenes containing people.

## 4. Discussion

While individuals with SCD and MCI have decreased memory performance in comparison to HC, there is a considerable overlap in the images that they remember and forget. Thus, there are images that are highly memorable and forgettable to everyone regardless of diagnosis. These consistencies in memorability exist not only between patient groups and healthy controls, where consistencies in memorability are already well-established for controls [1,2], but also within patient groups themselves. Our questionnaire-based assessment of image attributes revealed that this common memorability is not related to aesthetics or spaciousness, but to being manmade scenes that contain more objects, and are subjectively more memorable and interesting. While previous work has reported that ratings of interestingness, subjective memorability, and aesthetics are ultimately not predictive of scene memorability at a fine-grained scale for healthy populations [7], such attributes may be important for guiding the selection of images that are broadly memorable across population types.

Additionally, we show that a publicly available convolutional neural network (MemNet [6]) trained to predict image memorability also aligns with performance of HC as well as those with SCD and marginally with MCI. This raises the possibility that computational methods may guide the selection of images for diagnostic or therapeutic tools on the basis of memorability. Such tools may assist in creating or adapting environments to ease memory burdens on patients by avoiding low memorability items, or focusing strategies on rehearsing particularly forgettable information.

While memorability is generally consistent across HC, SCD, and MCI groups, we have also identified a specific set of images that significantly differ between groups. Namely, we find that there are images that are highly memorable to HC, yet highly forgettable to patients, and a certain subset of these images can be used to best determine if an individual is likely to be healthy or have MCI or SCD. The images generalize across impairments; images that differentiate MCI also successfully differentiate SCD, indicating that SCD may show similar cognitive impairments to those developed in MCI. This image set results in as much as a 10% improvement in diagnostic performance in comparison to a poorly chosen set of images (e.g., images memorable to patients but forgettable to healthy controls). Further, this optimized image set reaches peak diagnostic performance with as few as 18.3 images seen per participant, classifying as well as the original set with 88 images per participant. This means that individuals with MCI or SCD can be identified with higher certainty, and in a quicker, easier test. In terms of content, these diagnostic images tended to be manmade, indoor scenes that contained people. However, in contrast to memorable images, they tended to be less aesthetic, less interesting, and seem subjectively less memorable. Scenes containing people tend to be the most memorable [7], however it is perhaps the combination of memorable image content (e.g., people, manmade objects) yet lack of memorable qualities (e.g., interestingness, aesthetics) that causes these images to be remembered by healthy controls but forgotten by patients.

Functional neuroimaging work with healthy individuals has found that viewing memorable images results in automatic, stereotyped activity patterns in the visual cortex and medial temporal lobe [3,4]. In future work, investigating the neural fate of memorable and forgettable images in older individuals and those with SCD or MCI may aid in understanding how patients may differentially process images at different processing stages of perception and memory encoding. In the DELCODE study, we have indeed obtained fMRI data alongside the behavioral data reported here [11] and will be able to address this question in the future. A related question is how Alzheimer’s pathology is related to memorability. For instance, we have previously shown that increasing levels of CSF total-tau are related to decreasing novelty responses in the amygdala and the hippocampus [11]. These functional consequences of tau-pathology could influence memorability patterns in MCI or SCD. Indeed, activity in medial temporal lobe regions shows early and automatic sensitivity to the memorability of an image in healthy individuals [3]. Image diagnosticity as calculated in this study could also be related to the biomarker status of individuals, a possibility that we will be able to address in the future with larger sample sizes. It will also be important to better understand the features of an image that drive it to be forgettable, memorable, or diagnostic. While the current work uses a CNN trained on healthy participant memory data, as larger-scale patient data is collected, a CNN could learn to identify images that would be particularly effective in diagnosing patients.

In sum, we show the importance of images themselves in predicting what patients are likely to remember and differentiating patients from healthy individuals. Such insights will have a meaningful impact in how we design cognitive assessment tools and tests for early diagnosis of memory impairments, and in understanding how and why we process and remember certain images over others in our complex, visual world.

## Acknowledgements

The study was funded by the German Center for Neurodegenerative Diseases (Deutsches Zentrum für Neurodegenerative Erkrankungen (DZNE)), reference number BN012. W. Bainbridge is supported by the Intramural Research Program of the National Institutes of Health (ZIA-MH-002909), under National Institute of Mental Health Clinical Study Protocol 93-M-1070 (NCT00001360).

